# Intermittent suppressive posaconazole therapy is ineffective at mitigating cardiac and digestive tract pathologies in an experimental model of chronic Chagas disease

**DOI:** 10.1101/2024.12.03.626632

**Authors:** Shiromani Jayawardhana, Francisco Olmo, Amanda Fortes Francisco, Archie A. Khan, Michael D. Lewis, Martin C. Taylor, John M. Kelly

## Abstract

Infections with *Trypanosoma cruzi* cause Chagas disease, a chronic condition that can give rise to debilitating cardiac and/or gastrointestinal damage. However, it is unclear why only ∼30% of individuals progress to symptomatic pathology, why this can take decades to become apparent, and why there is such a wide range of disease outcomes. Disease pathology is a long-term cumulative process resulting from collateral damage caused by inflammatory immune responses that continually eliminate transient parasite infections in the heart and/or gastrointestinal tract. The guiding principle behind anti-parasitic drug development is that sterile cure is required to prevent progression to symptomatic pathology. Evidence suggests that the cumulative damage required to reach the symptomatic threshold is determined by a number of factors, including host and parasite genetics, which govern the intensity and location(s) of infection. Therefore, an alternative therapeutic strategy could involve long-term intermittent treatment, which may not confer sterile cure, but is able to suppress the parasite burden to a level where the disease does not become symptomatic within the life-time of the infected individual. To test this hypothesis, we used an experimental murine model that displays both cardiac and digestive tract pathology. Mice were given intermittent treatment with posaconazole, under conditions that reduced the parasite burden by >99%, but did not confer sterile clearance. Our results show that this did not provide long-term protection against the key cardiac or gastrointestinal manifestations of Chagas disease, and that sterile cure should remain the single goal of the drug development community.

## INTRODUCTION

Chagas disease, caused by the protozoan *Trypanosoma cruzi*, poses a public health challenge across Latin America (1). In addition, due to migration, it is increasingly detected in non-endemic regions (2, 3). Parasite transmission occurs primarily via triatomine bug vectors, but other routes of infection such as congenital transmission, consumption of contaminated food and drink, organ transplantation and blood transfusions also pose significant risk (4–7). Following *T. cruzi* infection, individuals enter the acute stage, which typically lasts 2-8 weeks, and is characterized by widespread parasite dissemination in blood and tissues. Symptoms at this stage are usually mild or non-specific, although outcomes can be fatal in some cases, due to myocarditis or encephalopathy. Immune responses eventually control, but do not eliminate the infection, and most individuals then remain asymptomatic for life (8–10). Approximately 30% of those infected ultimately develop chronic Chagas disease, although this can take decades to become symptomatic (11). *T. cruzi* infection results in chronic Chagas cardiomyopathy (CCC) in 20 - 30% of cases, and ∼10% lead to digestive Chagas disease (DCD), with the symptoms sometimes occurring in combination (12–14).

Several mechanisms have been postulated to explain the underlying causes of CCC, including direct parasite-mediated injury, parasite-induced inflammatory tissue damage, and autoimmune responses (15–18). The most widely accepted view is that pathology results from cumulative damage to heart tissue caused by pro-inflammatory immune responses driven by the presence of parasites. This in turn leads to fibrosis, cardiac hypertrophy and cardiomyopathy, although disease progression follows a complex pathway that can also involve arrhythmias and thromboembolisms, and is difficult to predict at an individual level (10, 19–21). During chronic *T. cruzi* infection, the parasite burden is extremely low and challenging to detect. In murine models, the gastrointestinal (GI) tract, skin, and in some strains, skeletal muscle, act as permissive reservoirs where infection foci persist (22–25). Cardiac infections in the chronic stage appear to be periodic rather than persistent, with detectable infection ranging from 10% - 80%, depending on the mouse-parasite strain combination. Persistent cardiac infection is not required for the development of fibrosis (22, 23). Rather, intermittent cardiac infection, perhaps a result of parasite trafficking from more immune-tolerant niches, seems to be sufficient. The resulting pro-inflammatory responses, continually eliminate these periodic infections, but often to the detriment of surrounding tissue (19). Cardiac muscle and nervous tissue have low regenerative capacity, a feature that may explain the progressive nature of the pathology (26).

In the case of DCD, outcomes can include changes in motility, secretion and absorption, particularly in the colon and oesophagus (14, 27, 28). Irreversible denervation in the enteric nervous system has been proposed to lead to progressive dysfunction in gut motility, involving difficulty in swallowing, slower GI transit and faeces retention, finally leading to megacolon or megaoesophagus (29, 30). Recently, a murine model of digestive Chagas disease has been developed in which a digestive transit delay and faeces retention is associated with chronic *T. cruzi* persistence in the colon (31). This model provides a platform for dissection of DCD pathogenesis (32).

For several decades, *T. cruzi* infections have been treated with the nitroheterocyclic drugs benznidazole and nifurtimox (33, 34). Despite their long-standing use, there are several issues which contribute to a high level of treatment failure (10-50%) (35, 36). These include treatment duration (60-90 days), toxicity that leads to patient non-compliance, and variable efficacy across the *T. cruzi* species. In addition, both these pro-drugs are bio-activated within the parasite by the same mitochondrial nitroreductase (TcNTR-1), which could lead to cross-resistance (37, 38). To address these challenges, there is an international effort, driven by large drug-development consortia, to extend the range of therapeutic options. A promising candidate that emerged early in this process was the anti-fungal drug posaconazole, an inhibitor of *T. cruzi* sterol 14α-demethylase (CYP51), an enzyme involved in ergosterol biosynthesis (39, 40). Posaconazole exhibited *in vitro* antiparasitic activity in the low nanomolar range, displayed promising *in vivo* efficacy, had a good safety profile, and was already licenced for human use (41, 42). Unfortunately, it ultimately failed in clinical trials (35, 43), and more sensitive *in vivo* studies in mice revealed an inability to provide regular sterile cure (44).

There is abundant evidence from animal experiments that curative benznidazole treatment can block the development of cardiac pathology, particularly if delivered early in an infection (17, 45, 46). Similarly, in the case of murine DCD, curative treatment in the acute stage halts disease progression, reverses the gut motility defect, and leads to a recovery in the density of colonic myenteric neurons (31, 32). In contrast, although non-curative treatment results an initial improvement in GI tract symptoms, this is transient, with infection relapse leading to a return of pathology. Collectively, these experiments provide a strong rationale for the consensus that the presence of the parasite is required to drive the immune-mediated tissue damage that results in Chagas disease pathology.

The Target Product Profile for new Chagas disease drugs describes a candidate that provides parasitological cure during the asymptomatic chronic stage of the infection (47). This is in accordance with our current understanding of disease pathogenesis, which predicts that timely intervention should block further development of pathology (17, 32). The discovery of new drugs that conform to this profile has been demanding (48). Given the progressive nature of Chagas disease, an alternative treatment strategy could involve long-term intermittent administration of therapeutics that are able to reduce the parasite burden sufficiently to slow the acquisition of tissue damage, so that symptomatic pathology and organ dysfunction does not arise during the life-time of the patient. To test this hypothesis, *T. cruzi* infected mice were treated long-term with posaconazole under conditions in which the parasite burden was greatly reduced, but not eliminated. The results strongly indicate that sterile cure is necessary to protect against the development of both cardiac and GI tract pathology.

## RESULTS

To determine if suppressive intermittent therapy could prevent or moderate the progression of Chagas disease pathology, posaconazole was selected because of its anti-parasitic profile (44) and low toxicity (41, 42). In addition, we had previously shown that this drug could reduce splenomegaly in mice chronically infected with *T. cruzi*, even when sterile cure was not achieved (44). We used female C3H/HeN mice infected with the bioluminescent *T. cruzi* JR-Luc strain (Materials and Methods). In this infection model, the acute stage parasite burden peaks 3 - 4 weeks post-infection (wpi) (23, 49), and mice develop chronic cardiac (17, 23, 50) and digestive (31, 32) pathology.

The experimental outline is shown in Fig. 1. Beginning 21 days post-infection (dpi), mice were treated with posaconazole, once daily for 12 days (20 mg/kg). By the end of this period, bioluminescence levels had been reduced by >99%, close to background levels (Fig. 2). One cohort of mice was given no further treatment (initial treatment group). The infection relapsed in all these mice, such that within 2 - 3 weeks of drug withdrawal, the parasite burden had returned to levels similar to those in non-treated control mice. There was then a reduction in parasite load, as the infection transitioned to the chronic stage (>11 wpi), with infection foci displaying a typically dynamic spatiotemporal profile (Fig. 2). After the initial treatment, a second cohort of mice was treated at monthly intervals (monthly treated group), for 5 days at 20 mg/kg, until the experimental end-point (31 wpi). With each mouse, monthly treatment resulted in a reduction in bioluminescence-inferred parasite burden to background levels, followed by a relapse. There was a tendency for the rebound in parasite burden to increase with time, such that later in the experimental schedule, it peaked at levels higher than in non-treated control mice (Fig. 2). In the third cohort, after the initial treatment period, mice were treated weekly with a single 20 mg/kg dose (weekly treatment group). With this regimen, parasites were barely detectable by *in vivo* imaging until 22 wpi, and as with the monthly dosing cohort, the extent of relapse then tended to increase, although this was not consistent (Fig. 2).

**FIG 1.**
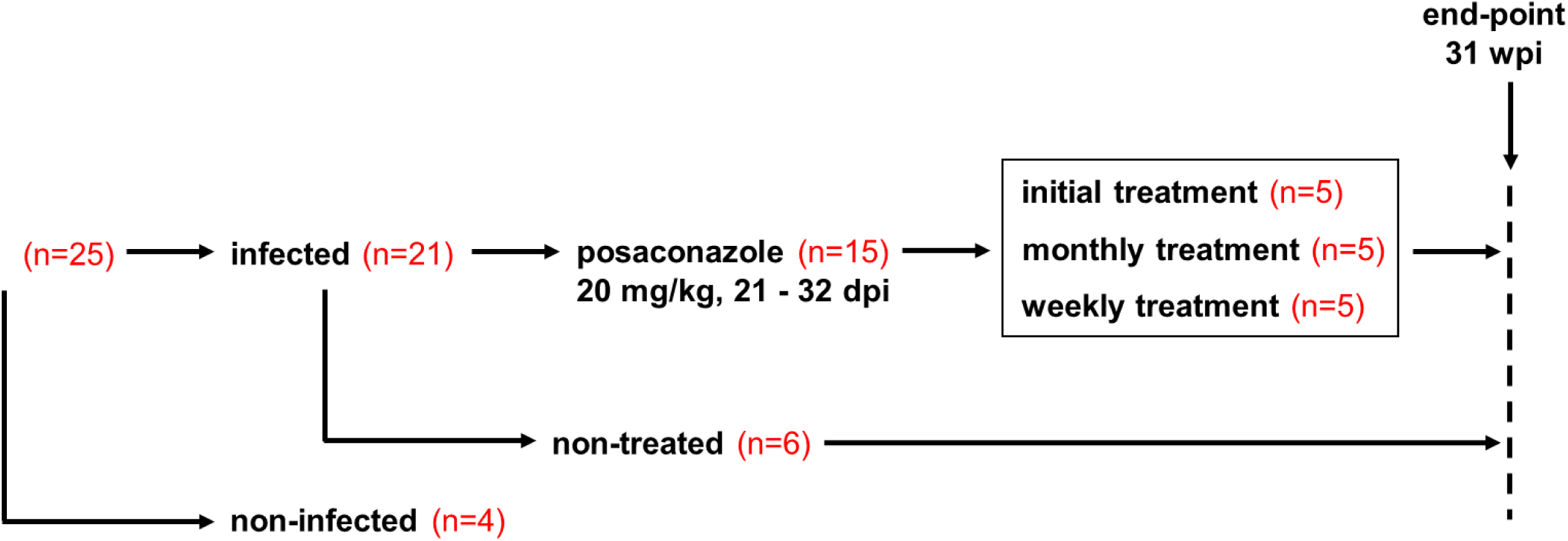
Experimental outline. C3H/HeN mice were infected with the bioluminescent. *T. cruzi* JR-Luc strain (DTU I). At 21 days post-infection (dpi), they were treated once daily with 20 mg/kg posaconazole for 12 days (Materials and Methods). One cohort of mice was given no further treatment. A second cohort was treated with posaconazole (20 mg/kg) for 5 days each month, and a third was treated one day each week until the experimental end-point at 31 weeks post-infection (wpi).

**FIG 2.**
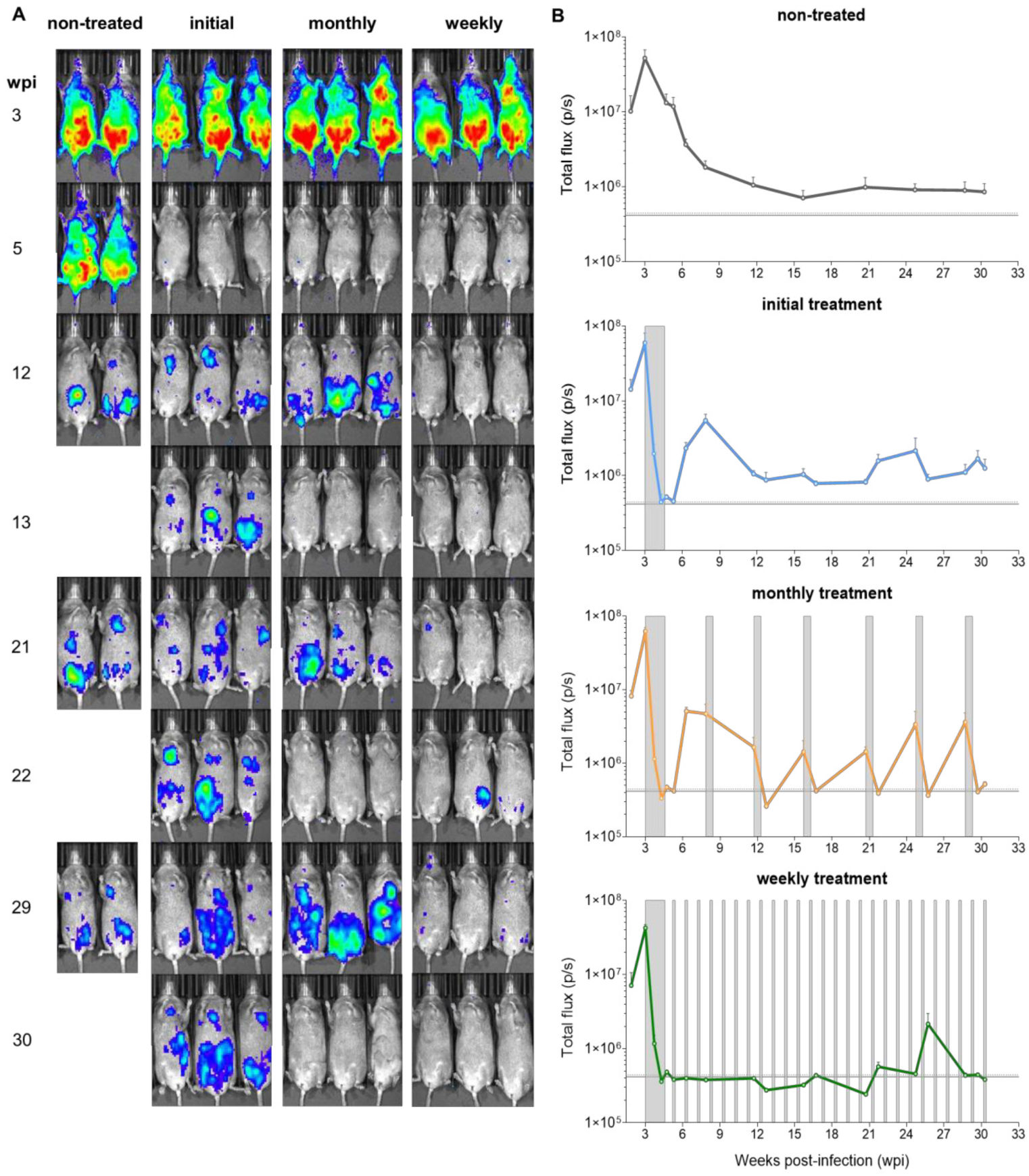
*In vivo* assessment of intermittent posaconazole treatment of *T. cruzi* infected mice. (A) C3H/HeN mice were infected with the bioluminescent *T. cruzi* JR-Luc strain and monitored regularly by *in vivo* imaging (ventral images shown). As outlined (Fig. 1), 21 dpi, mice were treated with posaconazole (20 mg/kg) once per day for 12 days. After this initial dosing, one group (n = 5) received no further treatment (initial). A second group (n = 5) was subsequently treated on 5 consecutive days each month (monthly), and a third group (n = 5) was treated weekly with a single dose of 20 mg/kg posaconazole (weekly). The heat-map is on a log10 scale and indicates intensity of bioluminescence from low (blue) to high (red). (B) Graphs showing the total body bioluminescence (ventral and dorsal imaging) (photons/second; p/s) of treated and non-treated mice throughout the experiment. The grey vertical bars indicate treatment periods.

Treated mice were examined by *ex vivo* imaging at the experimental end-point (Materials and Methods; Fig. 3). This confirmed that all mice in the treatment groups were still infected. Bioluminescent foci were detectable in the GI tract and skin in all cases, whereas in other organs and tissues, including the heart, infection was more sporadic. At the end-point, the overall level of infection in mice that received only the initial 12-day dosing, or received monthly intermittent treatment was similar to non-treated mice. Mice that received the single weekly dose had fewer detectable foci (Fig. 3), possibly reflecting that they had received their final posaconazole dose 3 days before the end-point at 31 wpi (Fig. 2). The limit of detection by *ex vivo* imaging is <12 parasites (24).

**FIG 3.**
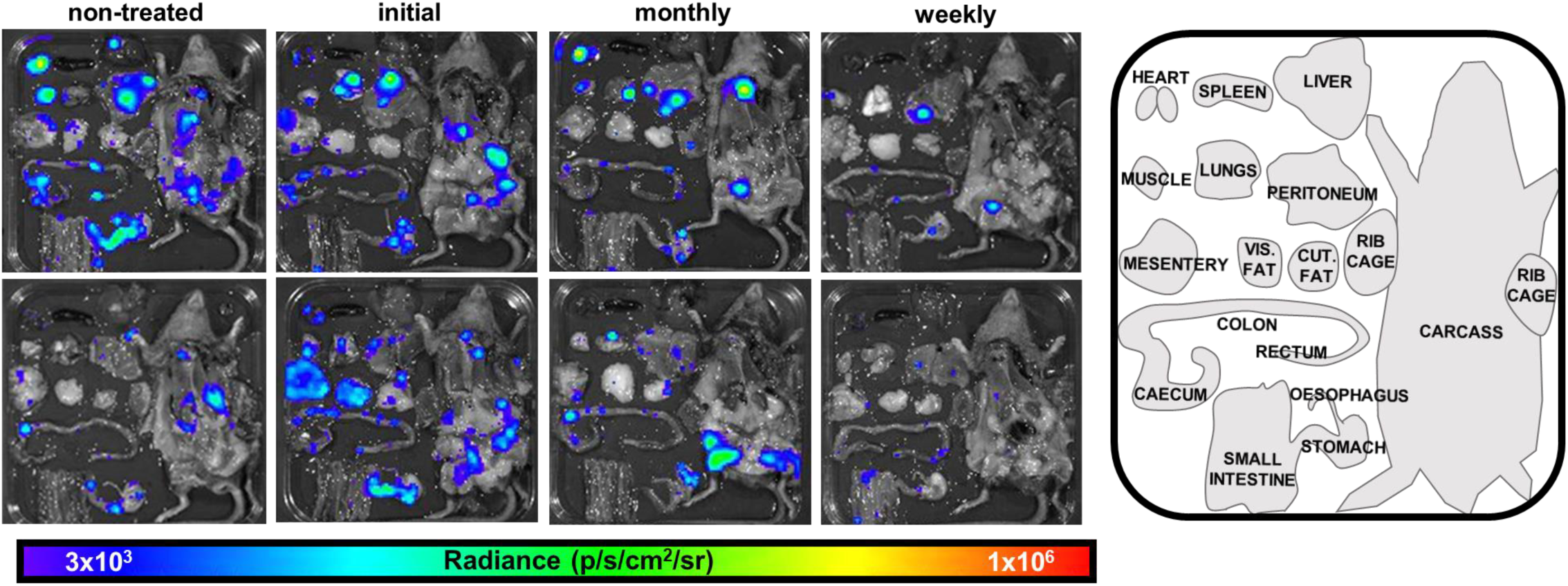
End-point tissue and organ distribution of *T. cruzi* following various posaconazole treatment regimens. (A) Two representative *ex vivo* images of tissues and organs harvested from mice in each cohort at the experimental end-point (31 wpi) (Materials and Methods). Tissues/organs were arranged as shown (inset, right). The heat-map is on a log10 scale and indicates intensity of bioluminescence from low (blue) to high (red).

To determine if the trend of relapses to become more pronounced over time was linked to acquired resistance (Fig. 2), 12 parasite clones were isolated from a range of mouse tissues (Materials and Methods). None of the clones displayed increased tolerance to posaconazole, as inferred from treatment of infected COLO-N680 cell cultures (Fig. S1).

Mice were assessed regularly throughout the experimental period for GI transit time delays as a marker for digestive Chagas disease after oral gavage with the red dye tracer carmine (Materials and Methods) (Fig. 4A, Fig. S2). As observed previously (31, 32), non-treated mice developed a delay during the acute stage, which peaked 4.5 wpi. After partially subsiding, severity increased later in the chronic stage (24 wpi and beyond) (Fig. 4A). Non-curative posaconazole treatment (initial treatment group) led to a rapid reversal of the GI transit delay, back to levels observed in the non-infected controls. However, the effect was temporary; the transit time reverted to that in non-treated mice by week 12, with severity increasing further by the experimental end-point (30 wpi). A detailed temporal breakdown and statistical analysis of the transit delay data are provided in Fig. S2. In the cohort of mice that were given additional monthly treatment, the reversal of the transit delay was maintained until beyond week 18, but then reached a level similar to that of non-treated mice by week 24 (Fig. 4A; Fig. S2). Treated mice given weekly doses of posaconazole, followed the same trend. At 30 wpi, several mice from all infected groups displayed transit time delays beyond the four-hour cut-off period, implying the development of severe chronic DCD pathology (Fig. 4A).

**FIG 4.**
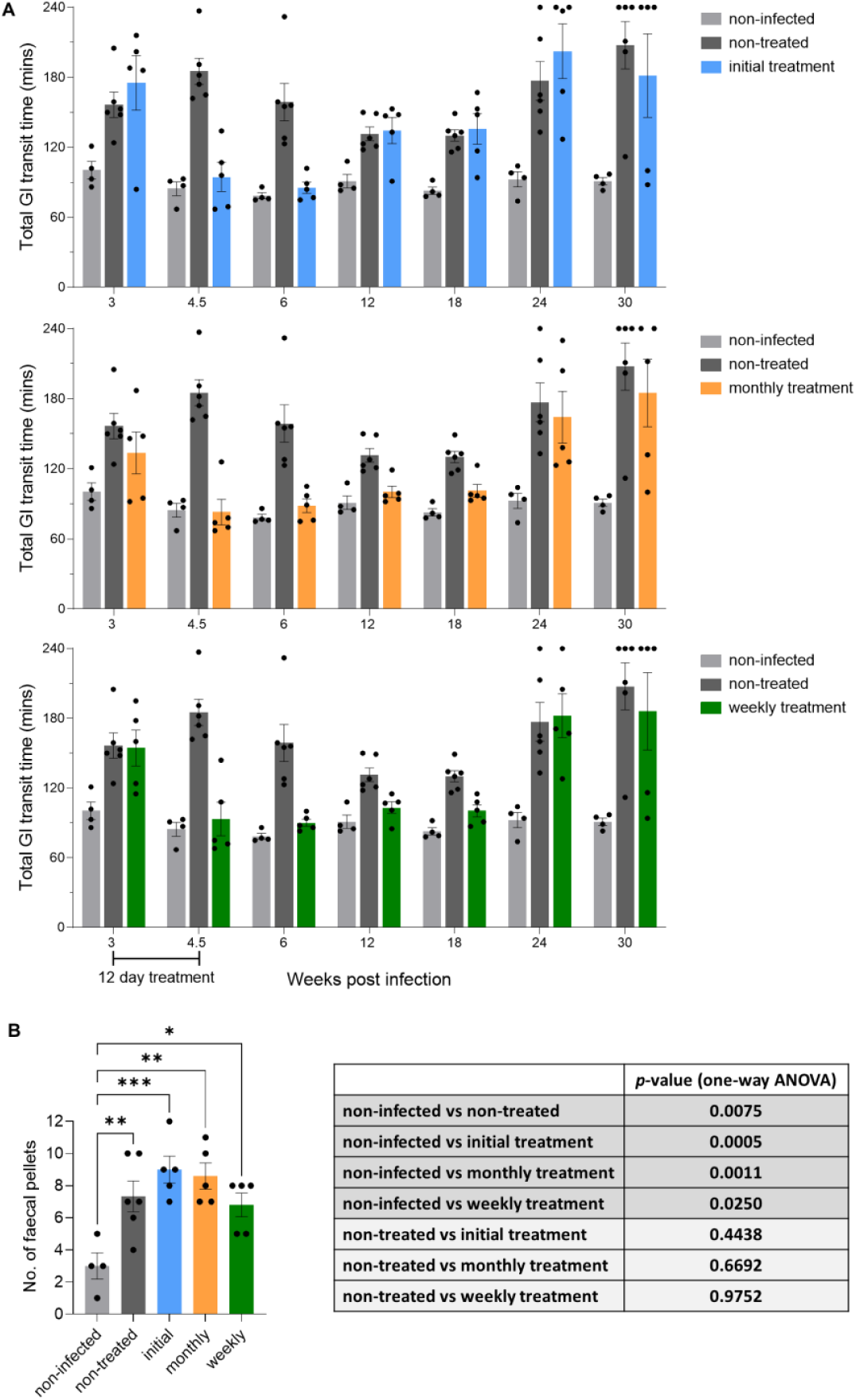
Intermittent posaconazole treatment does not protect *T. cruzi* infected mice from GI tract dysfunction. (A) Bar charts showing the impact of posaconazole treatment on GI transit time in C3H/HeN mice infected with the *T. cruzi* JR-Luc strain (non-infected mice, n=4; non-treated mice, n=6; each treatment group, n=5). Transit times were established by monitoring passage of the red dye tracer carmine (Materials and Methods). A 4-hour (240 minutes) cut-off point for transit data acquisition was imposed for animal welfare reasons. Each dot represents a single mouse. Data are expressed as mean ± SEM. See Fig. S1 for details on statistical analysis. (B) Bar chart showing the number of faecal pellets in the colon *post-mortem* (Materials and Methods) of non-infected mice, infected non-treated mice and infected mice treated with posaconazole (initial, monthly and weekly), as outlined in Fig. 1 and 2. Each dot represents a single mouse, with data expressed as mean ± SEM. Asterisks represent *p*-values for one-way ANOVA, followed by Dunnett’s multiple comparison post-hoc test (\**p*<0.05; \*\**p*<0.01; \*\*\**p*<0.001). All infected groups showed a significant difference to the non-infected group. There was no significant difference (*p*>0.05) between any of the treated groups and the non-treated group, as shown in the table.

At the experimental end-point, we also investigated the impact of the treatment regimens on the development of a constipation phenotype (31, 32) by assessing retention of faecal pellets in the colon after 2 hours fasting. Pellet numbers in each of the treated cohorts were found to be more than twice those in colons from non-infected mice (Fig. 4B), and not statistically different from mice that were non-treated. In this infection model, we previously observed that non-curative benznidazole treatment during the acute stage leads to a transient improvement in GI tract motility and function (32), but during the chronic stage, the extent of pathology reverts to that in non-treated mice. Our results here show that this chronic stage outcome also occurs even when suppressive intermittent treatment is administered at weekly or monthly intervals for the duration of the infection.

To evaluate the ability of suppressive posaconazole treatment to block development of cardiac pathology, quantitative analysis of collagen content was used as an indicator of pathological myocardial fibrosis (Materials and Methods). In non-infected age-matched control mice, the collagen content was notably consistent (15 cardiac images examined per mouse) (Fig. 5), and 3 - 4 times lower (*p*=0.01) than in non-treated infected mice. There was greater variation in the levels of fibrosis in infected mice, an observation that has been made previously (17), and which reflects the situation found in human infections. No significant difference in collagen content was observed between non-treated mice and those from the initial treatment group. Similarly, in the infected mice exposed to the weekly or monthly posaconazole treatment regimens, the extent of collagen deposition was not significantly different from the non-treated cohort, with no apparent benefit arising from the intermittent regimen.

**FIG 5.**
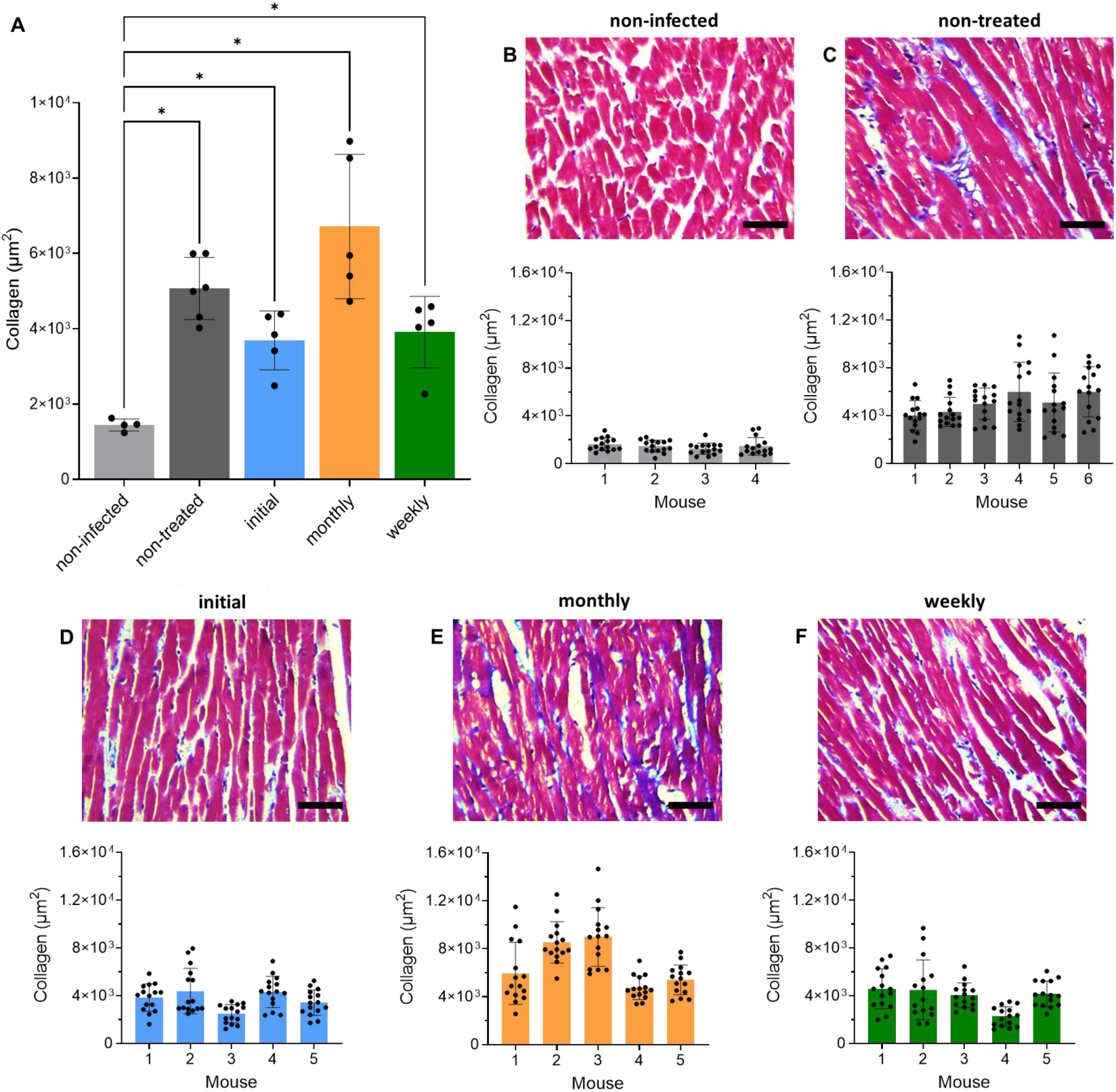
Intermittent posaconazole treatment does not protect *T. cruzi* infected mice from cardiac pathology. (A) Bar chart showing collagen content (blue area in Masson’s trichrome-stained sections) as a marker of cardiac fibrosis (Materials and Methods) in non-infected C3H/HeN mice (n=4), mice infected with *T. cruzi* JR-Luc strain, non-treated (n=6), and treated with posaconazole (n=5 per treatment cohort). Each dot corresponds to a single mouse, with data expressed as mean ± SEM. Asterisks represent *p*-values for one-way ANOVA, using Dunnett’s multiple comparison post-hoc test (\**p*<0.05), with a significant difference between the non-infected group and all infected groups. There was no significant difference (*p*>0.05) between any of the treatment groups and the non-treated group. (B-F) Representative Masson’s trichrome-stained photomicrographs highlighting collagen (blue) in cardiac sections from posaconazole-treated and non-treated mice. Magnification 400x, scale bars=100 µM. Bar charts show quantification of collagen content in individual mice from different groups, with each dot corresponding to the 15 randomly selected fields used for analysis, and data expressed as mean ± SEM.

## DISCUSSION

Chronic Chagas disease pathology is highly variable and can take decades to develop. Symptomatic pathology results from tissue damage caused by inflammatory immune responses to infections of the heart or GI tract, and accumulates over time (19, 32). The highly variable disease outcomes are thought to reflect the impact of host and parasite genetics, together with a range of environmental and life-style factors. Drug development strategies are based on a perceived need to achieve sterile cure (47). An alternative strategy could be the long-term use of drugs, which might not eliminate the infection, but is able to maintain the parasite burden below a threshold required to drive symptomatic pathology during the life-time of the infected individual. A crucial requirement of such drugs would be an acceptable safety profile under long-term use, for example, similar to that of statins.

To test this hypothesis, we selected posaconazole, a well-tolerated anti-fungal drug (41, 42) that reduces *T. cruzi* infections to very low levels, but typically does not eliminate the parasite (35, 43, 44). The initial treatment conditions (20 mg/kg for 12 days) reduced the parasite burden below the level of detection by *in vivo* imaging, but was followed by relapse in all cases (Fig. 2 and 3). Monthly treatment resulted in the parasite burden being temporarily reduced to background levels, only for it to rebound, sometimes higher than in the non-treated control mice, when drug pressure was released (Fig. 2). Despite these relapses, the continued ability of treatment to knockdown the parasite load suggested that long-term intermittent dosing did not result in the development of posaconazole-resistance. This was confirmed by assessing the posaconazole sensitivity of parasites rescued from infected mice at the experimental end-point (Fig. S1). Posaconazole is considered to be a trypanostatic drug, and activity against intracellular parasites *in vitro* reaches a plateau where increasing drug concentration no longer results in further growth inhibition (51). With *T. cruzi* JR-Luc, intracellular growth inhibition of the isolated parasite clones plateaued between 65-75% at all concentrations tested between 8 and 500 nM (Fig. S1). Increases in the extent of recrudescence after multiple suppressive treatments (Fig. 2A) might therefore reflect a selective bottleneck that allows parasites with a slightly higher *in vivo* growth rate to dominate the population.

DCD is an enteric neuropathy that in the C3H/HeN:TcJR-Luc host:parasite model results in a GI transit delay that first becomes apparent during the acute stage of infection (32). This is then followed by partial recovery, before further deterioration sets in as mice progress through the chronic stage. Drug-induced functional restoration following curative treatment with benznidazole in the acute stage is associated with recovery of neuronal density in the myenteric plexus (32). In the current study, we also observed a biphasic curve in GI transit time during the course of non-treated infections (Fig. 4A). Non-curative suppressive treatment with posaconazole, initiated during the acute stage (3 wpi), led to a rapid reversal of dysfunction, such that GI transit times returned to levels comparable with the non-infected control group. By week 12 however, the transit time in mice that had received only the initial posaconazole treatment returned to that in non-treated infected mice, and then underwent further deterioration (Fig. 4A). Both the monthly and weekly intermittent treatment groups exhibited reversal of the acute stage GI transit delay, and also displayed a lag in the onset of the chronic stage symptoms. However, by week 24, gut motility subsequently declined to levels exhibited in non-treated mice (Fig. 4A and Fig. S2). By the end-point, GI transit in the monthly and weekly intermittent treatment groups was significantly impaired in comparison with the non-infected group (Fig. S2). In benznidazole-cured mice, there is no deterioration in GI transit time during the chronic stage, indicating that a durable restoration of GI function is possible if a prompt sterile cure is achieved (32).

The partial improvement in GI transit times in non-treated mice towards the end of the acute stage, followed by further decline later in the chronic stage, suggests that two different mechanisms may be driving the GI transit pathology. This has parallels with CCC, where pathology in the acute stage is associated with myocarditis, and in the chronic stage, with cumulative damage and maladaptive fibrotic tissue repair that leads to cardiomyopathy (10). In the chronic stage, several mice in all the infected groups displayed GI transit time delays that were beyond the 4-hour cut-off (Fig. 4A), indicating the development of more severe DCD pathology than observed earlier in the infection. *T. cruzi* infection of the GI tract leads to cumulative neuron loss in the enteric nervous system, resulting in a progressive decline in gut motility, particularly associated with the colon (31). To further investigate the functional deterioration of peristalsis, end-point assessment of faecal retention in the colon was also carried out. This constipation phenotype, similar to that in human DCD, revealed no significant improvement in mice that received non-curative treatment, compared to non-treated controls. All the treated groups showed a significant increase in faecal retention (Fig. 4B). Therefore, under the conditions tested, long-term suppressive posaconazole treatment does not lead to improvement in chronic DCD symptoms, as judged by GI transit delay or faecal retention.

In C3H/HeN mice chronically infected with the *T. cruzi* JR-Luc strain, the profile of organ/tissue parasite distribution is more disseminated than in other murine models, and up to 80% of mice display cardiac-localised infections when assessed by *ex vivo* imaging (23, 44). Mice display extensive cardiac fibrosis, a marker of CCC (22), irrespective of whether their hearts are infected at the point of analysis. Consistent with the DCD data, our results show that long-term non-curative intermittent posaconazole treatment also provides no significant protection against cardiac fibrosis at the experimental end-point (Fig. 5A). Therefore, despite suppression of the overall parasite burden, particularly with the weekly treatment regimen, there was no significant reduction in the severity of either GI tract or cardiac pathology. One possibility might be that the transient and fluctuating nature of recrudescence could impact the dynamics of the inflammatory immune response, such that this negates any beneficial outcome arising from a reduction in the overall parasite burden.

In summary, we have examined the impact of two intermittent posaconazole treatment regimens on three key aspects of chronic Chagas disease pathology (GI transit delay, faecal retention and cardiac fibrosis). Our study demonstrates that these suppressive regimens do not prevent or alleviate long-term development of DCD or CCC, even when the parasite burden is considerably reduced. This outcome highlights that an ability to provide sterile cure should remain a pre-requisite of any anti-parasitic Chagas disease drug that is advanced to the clinic.

## MATERIALS AND METHODS

### Ethics statement

Animal studies were carried out under the UK Home Office licence P9AEE04E, in accordance with the UK Animals (Scientific Procedures) Act 1986, and approved by the London School of Hygiene and Tropical Medicine Animal Welfare and Ethical Review Board.

### Mouse husbandry, infections and treatment

The *T. cruzi* JR strain (Discrete Typing Unit I; DTU I), modified to express the red-shifted luciferase gene *PpyRE9h* (TcJR-Luc) (23, 52), was used in all experiments. Parasite growth and the generation of tissue culture trypomastigotes (TCTs) were as described previously (22). CB17-SCID mice purchased from Charles River (UK) were infected i.p. with 5×10^4^ TCTs in 0.2 ml PBS and used to derive blood trypomastigotes (BTs) via cardiac puncture. Female C3H/HeN mice were also purchased from Charles River (UK) and infected, aged 7-10 weeks, with 1×10^3^ BTs from SCID mice. Mice from each study group were housed in pathogen-free ventilated cages on a 12-hour light/dark cycle, with food and water *ad libitum*. Posaconazole (Biosynth Ltd; 20 mg/kg) was administered orally via gavage with a flexible cannula in a vehicle of 5% DMSO (v/v) and 95% (0.5% hydroxypropyl methylcellulose and 0.4% Tween 80 in Milli-Q H_2_O). At the 31-week experimental end-point, mice were euthanised by exsanguination under terminal anaesthesia (dolethal, 60 mg/kg i.p.) and necropsied, with organs excised and imaged for bioluminescence. Specific organs were collected for further analysis.

### *In vivo* and *ex vivo* bioluminescence imaging

Mice were injected with 150 mg/kg d-luciferin i.p. and anaesthetized after 5 minutes using 2.5% (v/v) gaseous isoflurane in oxygen (52, 53). Mice were then imaged after a further 5 minutes using the IVIS Spectrum system (Revvity, Hopkinton, MA, USA), with anaesthesia maintained through individual nose cones. Ventral and dorsal images were captured using Living Image v4.7.3, with exposure times of between 30 seconds and 5 minutes, depending on signal intensity. The threshold for *in vivo* imaging was set at 5 minutes exposure, large binning, using non-infected mice. Mice were then revived and returned to their cages.

To estimate parasite load during the course of infection, regions of interest (ROIs) using Living Image v4.7.3 were drawn around *in vivo* images of individual mice captured during bioluminescence imaging (53, 54). This was expressed as total flux (photons/second). Estimated detection threshold values for *in vivo* imaging were attained using ROI data from non-infected mice on set imaging days throughout the study.

Mice were fasted for 2 hours prior to necropsy for faecal pellet analysis. For *ex vivo* imaging, 150 mg/kg d-luciferin was i.p. injected, and mice were culled 5 minutes later by exsanguination under terminal anaesthesia. The mice were perfused with 10 ml 0.3 mg/ml d-luciferin in PBS, and organs excised and placed in a standardized arrangement on a Petri dish, along with the remaining carcass, and soaked in 0.3 mg/ml d-luciferin in PBS (53). Imaging of plated organs and carcass used 5 minutes exposure time and large binning.

### Parasite recovery from mouse tissue

Excised organs were maintained in ice-cold Hanks’ Balanced Salt Solution (HBSS) without calcium/magnesium, until disaggregation. Tissues were minced into 3-4 mm pieces using sterile scalpels, washed 3x with PBS to remove blood/debris, transferred to a tube containing a freshly made collagenase solution (100 U/mL collagenase IV in HBSS) and incubated at 37°C for 2 hours (55). The cell suspension was then filtered using a 200 µm cell strainer and centrifuged (300 g, 7 minutes). The pellet was resuspended in HBSS, and parasites were released using a 2 ml lysing matrix M (MP) in a Precellys 24 (Bertin) instrument, applying 2 cycles of 1 minute a 5000 speed. The suspension was then centrifuged (50 g, 5 minutes) to separate debris, and the supernatant was centrifuged (3000 g, 5 minutes) to pellet the parasites. Parasites were then resuspended in epimastigote medium (56) and cultured in 48-well plates in a CO_2_ incubator at 28°C.

### *In vitro* posaconazole sensitivity assay

To promote metacyclogenesis, TcJR-Luc epimastigotes were sub-cultured at 5×10^5^/ml and incubated for 14 days, until stationary phase (∼15% of the parasites transform). 1-2 ml from the culture surface was transferred to an Eppendorf tube, and swimming metacyclic trypomastigotes collected by centrifugation (3000 g, 5 minutes). The pellet was resuspended in Minimum Essential Medium (MEM), supplemented with 5% heat-inactivated foetal bovine serum (FBS), 100 U/ml penicillin, and 100 µg/ml streptomycin, and added to a 60-70% confluent culture of COLO-N680 cells (a human esophageal squamous cell carcinoma line). This was maintained at 37°C in a CO_2_ incubator with regular medium changes every 2-3 days. At day 8 post-infection, tissue-culture trypomastigotes (TCTs) in the cell supernatant were collected by centrifugation, resuspended in Dulbecco’s Modified Eagle Medium with high glucose (4 g/l), and counted.

Intrinsic luciferase expression in TcJR-Luc was used to monitor intracellular parasite replication. COLO-N680 cells (human oesophageal squamous cell carcinoma line) were seeded at 5×10^4^ cells per well in 100 µl growth medium in white, clear-bottomed 96-well microplates. After 8 hours, cells were infected overnight with 5×10^5^ TCTs/well. The following day, wells were washed 3x times with PBS to remove non-internalized TCTs, before adding 200 µl MEM supplemented with 2.5% FBS containing posaconazole at various concentrations. To assess activity, 8-point potency curves were generated by serial 2:1 dilution. At the experimental endpoint (96 hours), wells were washed 1x with PBS and incubated for 30 minutes in 100 µl fresh medium. Subsequently, 100 µl reconstituted Bright-Glo™ Luciferase Assay System reagent (Promega) was added, and plates were incubated at room temperature for 2 minutes in the dark to allow cell lysis. Bioluminescence was quantified in triplicate using a BMG FLUOstar Omega, with measurements obtained by integration over 3 seconds/well. Dose-response curves were fitted after calculating growth inhibition percentages compared to untreated controls for each clone. The 95% confidence intervals were calculated using the sigmoidal dose-response variable slope function in GraphPad Prism 10 software. Data are presented as the average of 2 independent experiments.

### GI transit time assay

200 µl 6% w/v carmine red dye solution in 0.5% methyl cellulose mixed in distilled water was administered to individual mice by oral gavage. Mice were then returned to their cages for 75 minutes, before being individually placed in separate containers. The time of the first red faecal pellet to be excreted was recorded. For animal welfare reasons, there was a maximum cut-off time of 4 hours. Time from gavage to excretion of the first red pellet was recorded as the total intestinal transit time. For faecal pellet analysis, colons were separated from other tissues and externally cleaned with PBS. The faecal pellets were eased out of the colon lumen and counted (31, 32).

### Histology and cardiac fibrosis

Mouse hearts were longitudinally bisected, placed in histology cassettes and fixed with Glyo-fixx for 24 hours at 4°C, then dehydrated in ethanol, cleared in xylene and embedded in paraffin blocks at 56°C (53). Heart sections (5 µm) were cut using a microtome, mounted on glass slides, dried overnight and stained with Masson’s trichrome. Quantification of collagen content was used as a marker for fibrosis, with 15 randomly selected fields per mouse used for analysis (17), with the investigator blinded to the groups. Using the binary processing prefilter function on the DFC420 light microscope (Leica Microsystems) and Leica Application Suite software v.4.2 for analysis, blue pixels were selected from the binary image for calculation of collagen fibres against the total area of cardiac muscle fibres staining red.

### Statistics

The unit of analysis was classified as individual mice, with no blinding or randomisation used. Statistical differences between groups were calculated using ordinary one-way ANOVA with Dunnett’s post-hoc correction for multiple comparisons, from GraphPad Prism v.8, with differences of *p*<0.05 considered significant.

---

**FIG S1. Posaconazole sensitivity of parasites isolated from mice at the experimental end-point.** (A) Tissue source of parasites isolated from each of the mouse groups. (B) 8-point dose-response curves were used to assess posaconazole sensitivity, based on luminescence (Materials and Methods). Growth inhibition plateaued at 65-75%, and remained unaltered up to 500 μM. (C) No significant differences in posaconazole sensitivity were found between parasites isolated from non-treated or treated mice.

**FIG S2. Statistical analysis of GI transit times between different groups.** Uninfected C3H/HeN mice (n=4) and C3H/HeN mice infected with the *T. cruzi* JR-Luc strain (non-treated, n=6; each treatment group, n=5). Each dot corresponds to a single mouse. Data of GI transit times are expressed as mean ± SEM, with all treated groups compared with non-treated and non-infected controls. A 4-hour (240 minutes) cut-off point for transit data acquisition was imposed for animal welfare reasons. Statistical analysis was carried out using ordinary one-way ANOVA, followed by Dunnett’s multiple comparison post-hoc test, with *p*-values of significant differences (*p*<0.05).

## Acknowledgements

This work was supported by UK Medical Research Council (MRC) grants MR/T015969/1 to J.M.K. and MR/R021430/1 to M.D.L., and funding from the Drugs for Neglected Diseases initiative (DNDi). DNDi received financial support from: Department for International Development (DFID), UK; Federal Ministry of Education and Research (BMBF) through KfW, Germany; and Médecins sans Frontières (MSF) International.

## REFERENCES

1. World Health Organisation. 2023. https://www.who.int/news-room/fact-sheets/detail/chagas-disease-(american-trypanosomiasis).

2. Irish A, Whitman JD, Clark EH, Marcus R, Bern C. 2022. Updated estimates and mapping for prevalence of Chagas disease among adults, United States. Emerg Infect Dis 28:1313–1320.

3. Gonzalez-Sanz M, Crespillo-Andújar C, Chamorro-Tojeiro S, Monge-Maillo B, Perez-Molina JA, Norman FF. 2023. Chagas disease in Europe. Trop Med Infect Dis 8:513.

4. de Fuentes-Vicente JA, Gutiérrez-Cabrera AE, Flores-Villegas AL, Lowenberger C, Benelli G, Salazar-Schettino PM, Córdoba-Aguilar A. 2018. What makes an effective Chagas disease vector? Factors underlying *Trypanosoma cruzi*-triatomine interactions. Acta Trop 183:23–31.

5. Edwards MS, Montgomery SP. 2021. Congenital Chagas disease: progress toward implementation of pregnancy-based screening. Curr Opin Infect Dis 34:538–545.

6. Robertson LJ, Havelaar AH, Keddy KH, Devleesschauwer B, Sripa B, Torgerson PR. 2024. The importance of estimating the burden of disease from foodborne transmission of *Trypanosoma cruzi*. PLoS Negl Trop Dis 18:e0011898.

7. Gómez LA, Gutierrez FRS, Peñuela OA. 2019. *Trypanosoma cruzi* infection in transfusion medicine. Hematol Transfus Cell Ther 41:262–267.

8. Rassi A Jr, Rassi A, Marin-Neto JA. 2010. Chagas disease. Lancet 375:1388–1402.

9. Pérez-Molina JA, Molina I. 2018. Chagas disease. Lancet 391:82–94.

10. Bonney KM, Luthringer DJ, Kim SA, Garg NJ, Engman DM. 2019. Pathology and pathogenesis of Chagas heart disease. Annu Rev Pathol 14:421–447.

11. Bonney KM, Engman DM. 2015. Autoimmune pathogenesis of Chagas heart disease: looking back, looking ahead. Am J Pathol 185:1537–1547.

12. Ribeiro AL, Nunes MP, Teixeira MM, Rocha MO. 2012. Diagnosis and management of Chagas disease and cardiomyopathy. Nat Rev Cardiol 9:576–589.

13. Cunha-Neto E, Chevillard C. 2014. Chagas disease cardiomyopathy: immunopathology and genetics. Mediat Inflamm 2014:683230.

14. Baldoni NR, de Oliveira-da Silva LC, Gonçalves ACO, Quintino ND, Ferreira AM, Bierrenbach AL, Padilha da Silva JL, Pereira Nunes MC, Ribeiro ALP, Oliveira CDL, Sabino EC, Cardoso CS. 2023. Gastrointestinal manifestations of Chagas disease: A systematic review with meta-analysis. Am J Trop Med Hyg 110:10–19.

15. Dc-Rubin SS, Schenkman S. 2012. *Trypanosoma cruzi* trans-sialidase as a multifunctional enzyme in Chagas disease. Cell Microbiol 14:1522–1530.

16. Tarleton RL, Zhang L, Downs MO. 1997. “Autoimmune rejection” of neonatal heart transplants in experimental Chagas disease is a parasite-specific response to infected host tissue. Proc Natl Acad Sci USA 94:3932–3937.

17. Francisco AF, Jayawardhana S, Taylor MC, Lewis MD, Kelly JM. 2018. Assessing the effectiveness of curative benznidazole treatment in preventing chronic cardiac pathology in experimental models of Chagas disease. Antimicrob Agents Chemother 62:e00832–18.

18. Kierszenbaum F. Chagas disease and the autoimmunity hypothesis. 1999. Clin Microbiol Rev 12:210–223.

19. Lewis MD, Kelly JM. 2016. Putting *Trypanosoma cruzi* dynamics at the heart of Chagas disease. Trends Parasitol 32:899–911.

20. Cutshaw MK, Sciaudone M, Bowman NM. 2023. Risk factors for progression to chronic Chagas cardiomyopathy: A systematic review and meta-analysis. Am J Trop Med Hyg 108:791–800.

21. Roman-Campos D, Marin-Neto JA, Santos-Miranda A, Kong N, D’Avila A, Rassi A Jr. 2024. Arrhythmogenic manifestations of Chagas disease: Perspectives from the bench to bedside. Circ Res 134:1379–1397.

22. Lewis MD, Fortes Francisco A, Taylor MC, Burrell-Saward H, McLatchie AP, Miles MA, Kelly JM. 2014. Bioluminescence imaging of chronic *Trypanosoma cruzi* infections reveals tissue-specific parasite dynamics and heart disease in the absence of locally persistent infection. Cell Microbiol 16:1285–1300.

23. Lewis MD, Francisco AF, Taylor MC, Jayawardhana S, Kelly JM. 2016. Host and parasite genetics shape a link between *Trypanosoma cruzi* infection dynamics and chronic cardiomyopathy. Cell Microbiol 18:1429–1443.

24. Ward AI, Lewis MD, Khan AA, McCann CJ, Francisco AF, Jayawardhana S, Taylor MC, Kelly JM. 2020. *In vivo* analysis of *Trypanosoma cruzi* persistence foci at single-cell resolution. mBio 11:e01242–20.

25. Pérez-Mazliah D, Ward AI, Lewis MD. 2021. Host-parasite dynamics in Chagas disease from systemic to hyper-local scales. Parasite Immunol 43:e12786.

26. Porrello ER, Mahmoud AI, Simpson E, Hill JA, Richardson JA, Olson EN, Sadek HA. 2011. Transient regenerative potential of the neonatal mouse heart. Science 331:1078–1080.

27. Pinazo MJ, Cañas E, Elizalde JI, García M, Gascón J, Gimeno F, Gomez J, Guhl F, Ortiz V, Posada Ede J, Puente S, Rezende J, Salas J, Saravia J, Torrico F, Torrus D, Treviño B. 2010. Diagnosis, management and treatment of chronic Chagas gastrointestinal disease in areas where *Trypanosoma cruzi* infection is not endemic. Gastroenterología y Hepatología 33:191–200.

28. Dantas RO. 2021. Management of esophageal dysphagia in Chagas disease. dysphagia 36:517–522.

29. de Oliveira RB, Troncon LE, Dantas RO, Menghelli UG. 1998. Gastrointestinal manifestations of Chagas disease. Am J Gastroenterol 93:884–889.

30. Arantes RM, Marche HH, Bahia MT, Cunha FQ, Rossi MA, Silva JS. 2004. Interferon-gamma-induced nitric oxide causes intrinsic intestinal denervation in *Trypanosoma cruzi*-infected mice. Am J Pathol 164:1361–1368.

31. Khan AA, Langston HC, Costa FC, Olmo F, Taylor MC, McCann, CJ, Kelly JM, Lewis MD. 2021. Local association of *Trypanosoma cruzi* chronic infection foci and enteric neuropathic lesions at the tissue micro-domain scale. PLoS Pathogens 17: e1009864.

32. Khan AA, Langston HC, Walsh L, Roscoe R, Jayawardhana S, Francisco AF, Taylor MC, McCann CJ, Kelly JM, Lewis MD. 2024. Enteric nervous system regeneration and functional cure of experimental digestive Chagas disease with trypanocidal chemotherapy. Nature Comm 15:4400.

33. Wilkinson SR, Kelly JM. 2009. Trypanocidal drugs: mechanisms, resistance and new targets. Exp Rev Molec Med 11:e31, p1–24.

34. Lascano F, García Bournissen F, Altcheh J. 2022. Review of pharmacological options for the treatment of Chagas disease. Br J Clin Pharmacol 88:383–402.

35. Molina I, Gómez i Prat J, Salvador F, Treviño B, Sulleiro E, Serre N, Pou D, Roure S, Cabezos J, Valerio L, Blanco-Grau A, Sánchez-Montalvá A, Vidal X, Pahissa A. 2014. Randomized trial of posaconazole and benznidazole for chronic Chagas disease. N Engl J Med 370:1899–1908.

36. Morillo CA, Marin-Neto JA, Avezum A, Sosa-Estani S, Rassi A Jr, Rosas F, Villena E, Quiroz R, Bonilla R, Britto C, Guhl F, Velazquez E, Bonilla L, Meeks B, Rao-Melacini P, Pogue J, Mattos A, Lazdins J, Rassi A, Connolly SJ, Yusuf S; BENEFIT Investigators. 2015. Randomized trial of benznidazole for chronic Chagas cardiomyopathy. N Engl J Med 373:1295–1306.

37. Wilkinson SR, Taylor MC, Horn D, Kelly JM, Cheeseman I. 2008. A mechanism for cross-resistance to nifurtimox and benznidazole in trypanosomes. Proc Natl Acad Sci USA 105:5022–5027.

38. Mejia AM, Hall BS, Taylor MC, Gómez-Palacio A, Wilkinson SR, Triana-Chávez O, Kelly JM. 2012. Benznidazole-resistance in Trypanosoma cruzi is a readily acquired trait that can arise independently in a single population J Inf Dis 206:220–228.

39. Urbina JA. 2009. Ergosterol biosynthesis and drug development for Chagas disease. Mem Inst Oswaldo Cruz 104:Suppl 1, 311–318.

40. Lepesheva GI, Villalta F, Waterman MR. 2012. Targeting *Trypanosoma cruzi* sterol 14α-demethylase (CYP51). Adv Parasitol 75:65–87.

41. Raad II, Graybill JR, Bustamante AB, Cornely OA, Gaona-Flores V, Afif C, Graham DR, Greenberg RN, Hadley S, Langston A, Negroni R, Perfect JR, Pitisuttithum P, Restrepo A, Schiller G, Pedicone L, Ullmann AJ. 2006. Safety of long-term oral posaconazole use in the treatment of refractory invasive fungal infections. Clin Infect Dis 42:1726–1734.

42. Wong TY, Loo YS, Veettil SK, Wong PS, Divya G, Ching SM, Menon RK. 2020. Efficacy and safety of posaconazole for the prevention of invasive fungal infections in immunocompromised patients: a systematic review with meta-analysis and trial sequential analysis. Sci Rep 10:14575.

43. Morillo CA, Waskin H, Sosa-Estani S, Del Carmen Bangher M, Cuneo C, Milesi R, Mallagray M, Apt W, Beloscar J, Gascon J, Molina I, Echeverria LE, Colombo H, Perez-Molina JA, Wyss F, Meeks B, Bonilla L.R, Gao P, Wei B, McCarthy M, Yusuf S, 2017. Benznidazole and posaconazole in eliminating parasites in asymptomatic *T. cruzi* carriers: The STOP-CHAGAS trial. J Amer Coll Cardiol 69:939–947.

44. Fortes Francisco A, Lewis MD, Jayawardhana S, Taylor MC, Chatelain E, Kelly JM. 2015. The limited ability of posaconazole to cure both acute and chronic *Trypanosoma cruzi* infections revealed by highly sensitive *in vivo* imaging. Antimicrob Agents Chemother 59:4653–4661.

45. Santos FM, Mazzeti AL, Caldas S, Gonçalves KR, Lima WG, Torres RM, Bahia MT. 2016. Chagas cardiomyopathy: The potential effect of benznidazole treatment on diastolic dysfunction and cardiac damage in dogs chronically infected with *Trypanosoma cruzi*. Acta Trop 161:44–54.

46. Calvet CM, Choi JY, Thomas D, Suzuki B, Hirata K, Lostracco-Johnson S, de Mesquita LB, Nogueira A, Meuser-Batista M, Silva TA, Siqueira-Neto JL, Roush WR, de Souza Pereira MC, McKerrow JH, Podust LM. 2017. 4-aminopyridyl-based lead compounds targeting CYP51 prevent spontaneous parasite relapse in a chronic model and improve cardiac pathology in an acute model of *Trypanosoma cruzi* infection. PLoS Negl Trop Dis 11:e0006132.

47. Drugs for Neglected Diseases initiative. 2024. https://dndi.org/diseases/chagas/

48. Francisco AF, Jayawardhana S, Olmo F, Lewis MD, Wilkinson SR, Taylor MC, Kelly JM. 2020. Challenges in Chagas disease drug development. Molecules 25: 2799.

49. Olmo F, Jayawardhana S, Khan AA, Langston HC, Francisco AF, Atherton RL, Ward AI, Taylor MC, Kelly JM, Lewis MD. 2024. A panel of phenotypically and genotypically diverse bioluminescent:fluorescent *Trypanosoma cruzi* strains as a resource for Chagas disease research. PLoS Negl Trop Dis 18:e0012106.

50. Francisco AF, Sousa GR, Vaughan M, Langston H, Khan A, Jayawardhana S, Taylor MC, Lewis MD, Kelly JM. 2023. Cardiac abnormalities in a predictive mouse model of Chagas disease. Pathogens 12:1364.

51. Moraes CB, Giardini MA, Kim H, Franco CH, Araujo-Junior AM, Schenkman S, Chatelain E, Freitas-Junior LH. 2014. Nitroheterocyclic compounds are more efficacious than CYP51 inhibitors against *Trypanosoma cruzi*: implications for Chagas disease drug discovery and development. Sci Rep 4:4703.

52. Branchini BR, Ablamsky DM, Davis AL, Southworth TL, Butler B, Fan F, Jathoul AP, Pule MA. 2010. Red-emitting luciferases for bioluminescence reporter and imaging applications. Anal Biochem 396:290–297.

53. Taylor MC, Francisco AF, Jayawardhana S, Mann GS, Ward A, Olmo F, Lewis MD, Kelly JM. 2019. Exploiting genetically modified dual-reporter strains to monitor experimental *Trypanosoma cruzi* infections and host:parasite interactions. Karina Andrea Gomez and Carlos Andres Buscaglia (eds.), T. cruzi Infection: Methods and Protocols, Methods Molec Biol 1955:147-163.

54. Lewis MD, Fortes Francisco A, Taylor MC, Kelly JM. 2015. A new experimental model for assessing drug efficacy against *Trypanosoma cruzi* infection based on highly sensitive *in vivo* imaging. J Biomolec Screening 20:36–43.

55. Freshney R. 1987. Culture of animal cells: A manual of basic techniques, p117, Alan R. Liss Inc., New York.

56. Kendall G, Wilderspin AF, Ashall F, Miles MA, Kelly JM. 1990. *Trypanosoma cruzi* glycosomal glyceraldehyde-3-phosphate dehydrogenase does not conform to the ‘hotspot’ topogenic signal model. EMBO J 9:2751–2758.

